# Memory deficits in males and females long after subchronic immune challenge

**DOI:** 10.1101/379339

**Authors:** Daria Tchessalova, Natalie C. Tronson

## Abstract

Memory impairments and cognitive decline persist long after recovery from major illness or injury, and correlate with increased risk of later dementia. Here we developed a subchronic peripheral immune challenge model to examine delayed and persistent memory impairments in females and in males. We show that intermittent injections of either lipopolysaccharide or Poly I:C cause memory decline in both sexes that are evident eight weeks after the immune challenge. Importantly, we observed sex-specific patterns of deficits. Females showed impairments in object recognition one week after challenge that persisted for at least eight weeks. In contrast, males had intact memory one week after the immune challenge but exhibited broad impairments in memory tasks including object recognition, and both context and tone fear conditioning several months later. The differential patterns of memory deficits in males and in females were observed without sustained microglial activation or changes in blood-brain barrier permeability. Together, these data suggest that transient neuroimmune activity results in differential vulnerabilities of females and males to memory decline after immune challenge. This model will be an important tool for determining the mechanisms in both sexes that contribute to memory impairments that develop over the weeks and months after recovery from illness. Future studies using this model will provide new insights into the role of chronic inflammation in the pathogenesis of long-lasting memory decline and dementias.

## 1. INTRODUCTION

Long-lasting memory dysfunction is common after a critical illness or major surgery. More than 25% of patients develop memory impairments that persist for months to years after recovery from major illness (Gharacholou et al., 2011; Semmler et al., 2013). These memory deficits can progress over time, contributing to neurodegenerative disorders and dementias including Alzheimer’s disease (Evered et al., 2016). Chronic low-grade inflammatory signaling is also correlated with memory decline (Noble et al., 2009, 2010; Yaffe et al., 2003) and has been implicated as both a cause and a consequence of dementia (Stephenson et al, 2018). In this study, we examine the causal role of a subchronic peripheral immune challenge in progressive memory decline.

Immune signaling is critically important for normal neural function, and activation of the peripheral immune system modulates memory and other cognitive processes (Marin & Kipnis, 2017; Wilson, Finch, & Cohen, 2002; Yirmiya & Goshen, 2011). In animal models, systemic immune activation results in memory deficits during acute inflammatory signaling (Czerniawski & Guzowski, 2014; Kranjac et al., 2012; Pugh et al., 1998). Hippocampal-dependent memory tasks including novel object recognition, spatial memory, and context fear conditioning are all sensitive to disruption by an acute systemic inflammatory challenge (Frühauf et al., 2015; Kahn et al., 2012; Pugh et al., 1998) suggesting that the hippocampus is particularly susceptible to dysregulation by immune activation (Williamson & Bilbo, 2013).

Previous work has demonstrated the long-lasting consequences of an overwhelming immune challenge on neural and cognitive function and for neuroimmune signaling. The cecal ligation and puncture model of sepsis results in impairments of memory that emerge soon after surgery (e.g., Barichello et al., 2007) and persist for months (e.g., Huerta et al., 2016). Similarly, a single high-dose injection of lipopolysaccharide results in memory deficits at least 5 months post-injection (Ming, Sawicki, & Bekar, 2015). Ongoing neuroimmune signaling, specifically microglial activation, in the weeks and months after an overwhelming inflammatory insult has been associated with memory deficits (Giustina et al., 2017; Semmler et al., 2007; Weberpals et al., 2009). In addition, persistent changes in blood brain barrier (Erickson & Banks, 2018) are commonly observed after illness (Sakusic & Rabinstein, 2018) and injury (Prakash & Carmichael, 2017) and proposed as a mechanism contributing to persistent neuroimmune changes and a variety of neurological and psychiatric disorders (Saunders, Ek, Habgood, & Dziegielewska, 2008). Nevertheless, since disease processes persist long after the initial immune challenge in these models, it is not clear whether sustained neuroimmune dysregulation or immune-triggered changes in neural processes mediates memory deficits long after a systemic immune challenge.

Lower intensity immune challenges may be more relevant to modeling progressive memory impairments as a consequence of chronic, low-grade inflammation (e.g., Yaffe et al., 2003). Repeated, lower dose immune challenges also cause impairments of memory (Kahn et al., 2012; Weintraub et al., 2013, 2014) and neural plasticity (Maggio, Shavit-Stein, Dori, Blatt, & Chapman, 2013) in the first weeks after injection. In this project, we determined whether a moderate, subchronic immune challenge results in persistent memory deficits months after immune challenge, whether memory deficits are associated with lasting neuroimmune activity, and the susceptibility of males and females to immune-mediated memory deficits.

Given the greater vulnerability of women to memory disorders such as Alzheimer’s disease (Snyder et al., 2016), we are particularly interested in the differential impact of subchronic immune challenge on males and females. Sex differences in peripheral responses are well described (Ghosh & Klein, 2017; Scotland, Stables, Madalli, Watson, & Gilroy, 2011) and there is growing evidence for sex differences in neuroimmune responses and function (Estefanía Acaz-Fonseca, Avila-Rodriguez, Garcia-Segura, & Barreto, 2016; Engler et al., 2016; Sorge et al., 2016; Speirs & Tronson, 2018). It is likely, therefore, that males and females are differentially susceptible to memory decline after subchronic immune challenge. To date, all studies of the lasting consequences of immune challenge on memory have been conducted with male animals.

Here we identified a causal role for subchronic immune activation on memory deficits in the months following recovery in both sexes. We demonstrated that males show delayed memory deficits in hippocampal-and amygdala-dependent tasks that emerged several months after immune challenge. Females showed impaired object recognition memory soon after immune challenge and these deficits persisted for at least eight weeks. These memory deficits were evident despite no evidence for long-lasting microglial activation or blood-brain barrier permeability.

## 2. MATERIALS AND METHODS

### 2.1 Animals.

9-11 week old male and female C57BL/6N mice from Envigo (Indianapolis, IN) were used in all experiments. Mice were individually housed with mouse chow and water provided *ad libitum* as previously described (Keiser et al., 2017). Individual housing in mice prevents fighting-induced stress (Meakin et al., 2013) and is ethologically appropriate for males and females (Becker & Koob, 2016). Individual housing is suitable for testing novel object recognition (Vogel-Ciernia & Wood, 2015) and contextual fear conditioning (Keiser et al., 2017) and follows the University of Michigan Institutional Care and use Committee policies on managing fighting in mice. The facility is ventilated with constant air exchange (60 m^3^/ h), temperature (22 ±1°C), and humidity (55±10%) with a standard 12 h light-dark cycle. Experiments were performed during the light portion of the cycle. Mice were acclimated to the colony room for at least seven days prior to injections. All experimental methods used in these studies were approved by the University of Michigan Committee on the Use and Care of Animals.

### 2.2 Immune stimulants.

Lipopolysaccharide (LPS, Escherichia coli, serotype 0111:B4; Sigma-Aldrich, St. Louis) was dissolved in saline (12.5μg/mL) and was injected intraperitoneally (i.p.; 250μg/kg; Fruhauf et al., 2015). Polyinosinic:polycytidylic acid (Poly I:C, P9582; Sigma-Aldrich) lyophilized powder was dissolved in distilled deionized water (10mg/ml), heated to 50°C and cooled to allow re-annealing, and injected i.p. (6mg/kg; Cunningham, Campion, Teeling, Felton, & Perry, 2007). These doses of LPS and Poly I:C were chosen based on previous reports of efficacy of these doses on memory-related paradigms (e.g., (Frühauf et al., 2015; Kranjac et al., 2011), for their similar and transient effects on eight loss (Fig. 1), and for their minimal effect on observed sickness behaviors in our laboratory in either sex.

**Figure 1.**
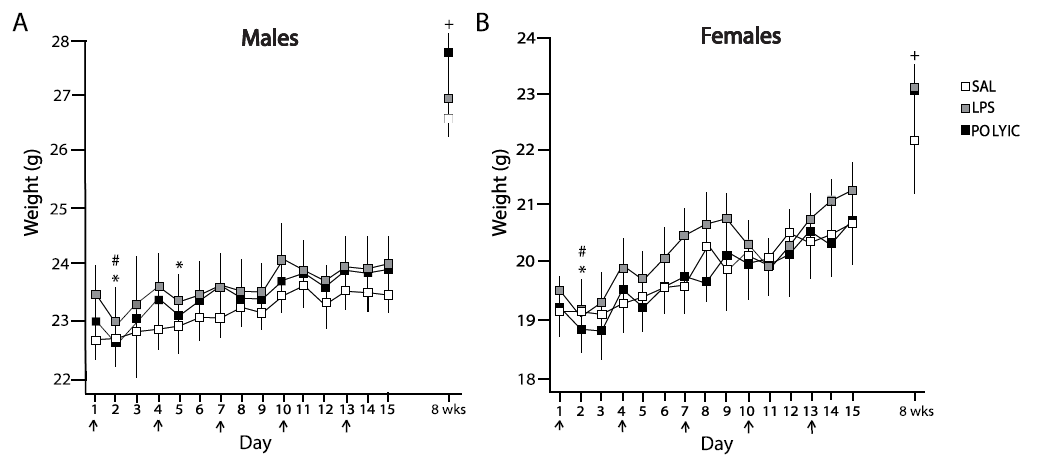
Experimental design and mouse weights after immune challenge. (A) Males and (B) females showed weight loss after initial LPS and Poly I:C injections, but showed no persistent changes in weight months after immune challenge. Arrows indicate injection days. SAL: saline; LPS: lipopolysaccharide; Poly I:C: Polyinosinic:polycytidylic acid. \**p* < 0.05 compared with injection day LPS group; #*p* < 0.05 compared with injection day Poly I:C group, +*p* < 0.05 Poly I:C vs Saline.

### 2.3 Subchronic Immune Challenge.

Mice received five intermittent injections of LPS (250μg/kg; n= 8-9), Poly I:C (6 mg/kg; n= 8-9), or saline control (n = 8-9), spaced three days apart. All injections were performed at the same time of day (Roberts, 2000). Mice were weighed daily throughout the injection period and weekly until testing. Changes in weight were assessed using a repeated-measures (Day × ImmuneChallenge) ANOVA.

### 2.4 Behavioral Testing.

Memory tests began one (Figs. 2-3) or eight weeks (Figs. 4-7) after the final injection. All testing was completed within 12 weeks after subchronic immune challenge. In experiments in which animals were tested on multiple tasks (e.g., Figs. 4 & 6; Figs. 5B-D & 7B-E), novel object recognition (3-8 days of testing) was always conducted first, followed by context fear conditioning (3 days of testing), with 3-10 days between tasks.

**Figure 2.**
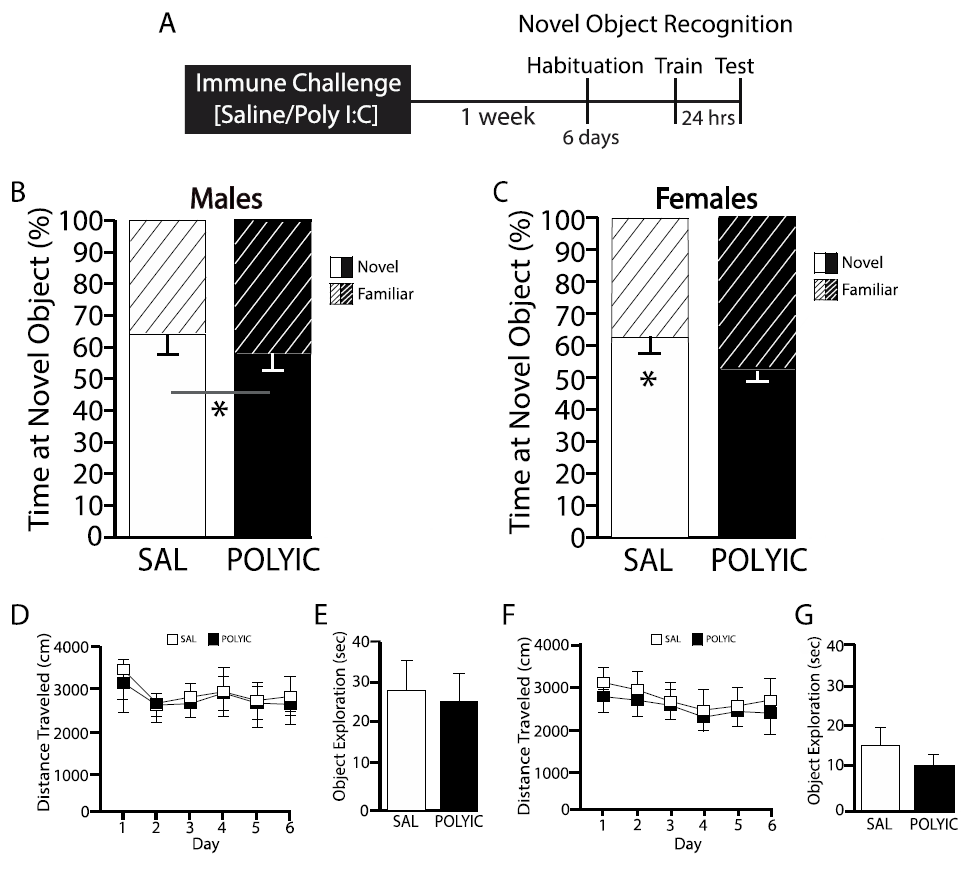
Hippocampal-dependent novel object recognition memory was disrupted only in females soon after subchronic Poly I:C. (A) Experimental timeline. Novel object recognition consisted of 6 days of habituation starting 8 weeks after last injection, training the day after habituation. Tests occurred 24 hours after training. (B) Intact novel object recognition in males shortly after Poly I:C (n = 8). (C) Subchronic Poly I:C treatment disrupted long-term novel object recognition memory in females in the weeks soon after the final injection (n = 8). (D, F) Neither habituation nor locomotor activity were altered one week after Poly I:C. (E, G) Total object exploration was similar amongst experimental groups during training in males and in females. \**p* <

**Figure 3.**
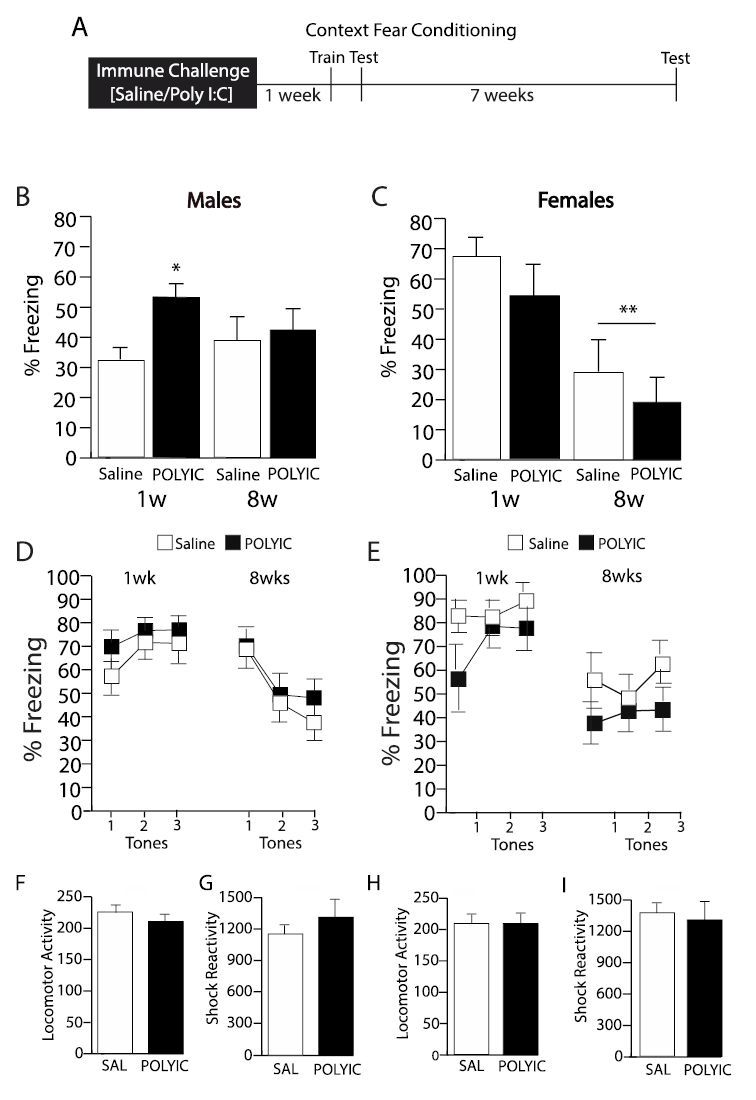
Subchronic Poly I:C did not cause early context fear memory impairments or retrieval deficits. (A) Experimental timeline. Animals were trained in context fear conditioning one week after the final injection. Context fear conditioning was assessed 24 hours, and auditory‐ cued fear test 48 hours after training. Animals were tested for fear memory retrieval seven weeks after training. (B) Prior Poly I:C challenge did not disrupt context fear conditioning soon after immune challenge, or during memory retrieval seven weeks later in males (n = 7-8). (C) In females, no disruption of context fear conditioning was observed soon after immune challenge; and, all females showed decreased freezing at the remote test, with no effect of prior immune challenge. (D,E) Tone fear conditioning remained intact soon after subchronic immune challenge in both sexes, and retrieval remained intact seven weeks later. (F,H) Locomotor activity and (G,I) shock reactivity were equivalent across all groups. \**p* < 0.05 *cf* Saline; #*p* < 0.001 *cf* 1week.

**Figure 4.**
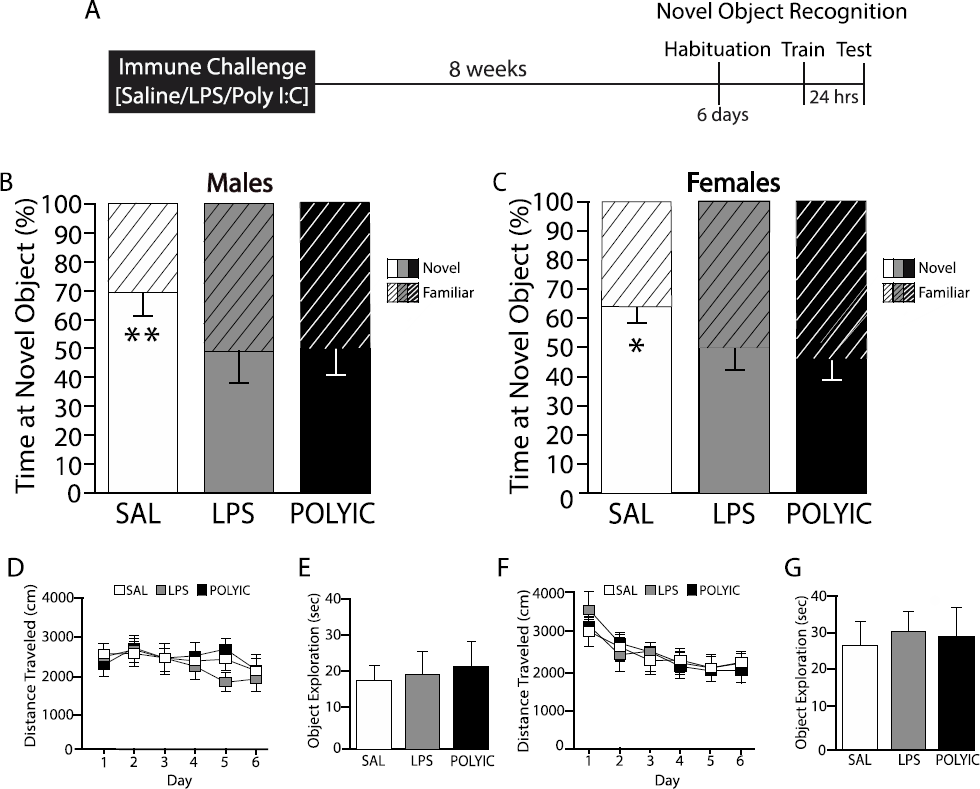
Impairments of hippocampal-dependent novel object recognition memory persisted long after subchronic immune challenge. (A) Experimental timeline. Habituation (6 days) began 8 weeks after last injection, training the day after habituation, and testing 24 hours after training. (B,C) Saline-treated males and females spent more time exploring the novel object (solid bar) compared with the familiar object (striped bar) 24 hours after training. Long-term memory for novel object recognition was disrupted in LPS‐ and Poly I:C treated mice (n = 8-9 animals/ group). (D,F) Neither locomotor activity nor habitation were affected by prior immune challenge. (E,G) There were no differences in total object exploration during training. \**p* < 0.05, \*\**p* <

**Figure 5.**
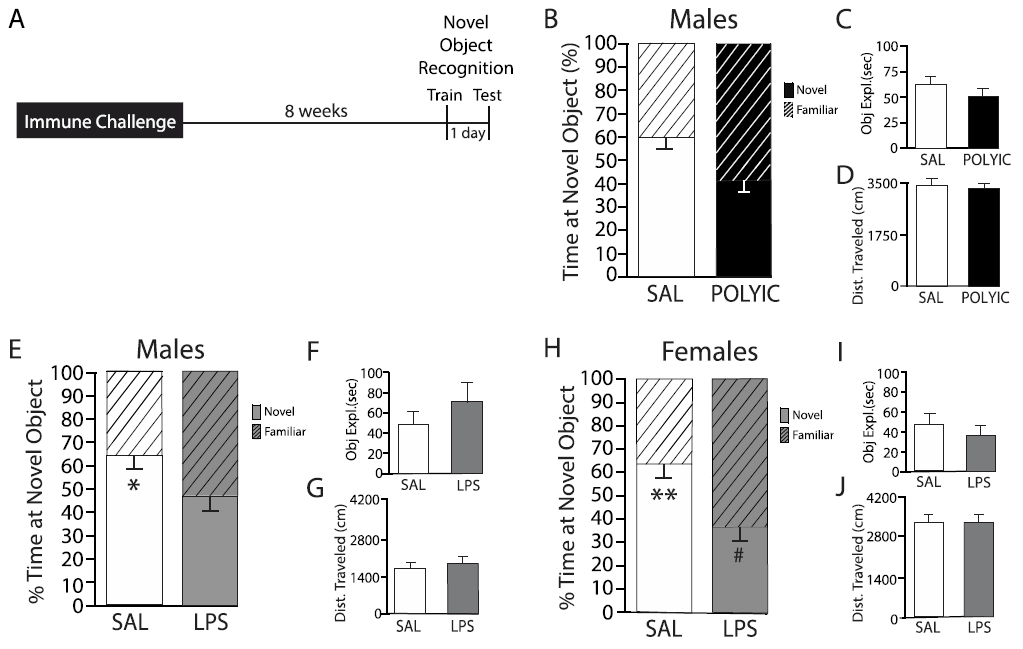
Subchronic immune challenge caused impaired 3-hour novel object recognition long after subchronic Poly I:C or LPS challenge. (A) Experimental timeline, with novel object recognition training and testing 8 weeks after last injection. (B) Saline-treated mice showed preference for the novel object when tested 3 hours after training, and this was disrupted months after Poly I:C. (C,D) Total object exploration and locomotor activity during novel object training were similar between groups. (E,H) Short-term memory for novel object recognition was impaired in males and females when tested 3 hours after training. (G,I) Total exploration during training and (H,J) locomotor activity were not altered by prior LPS in either sex.

**Figure 6.**
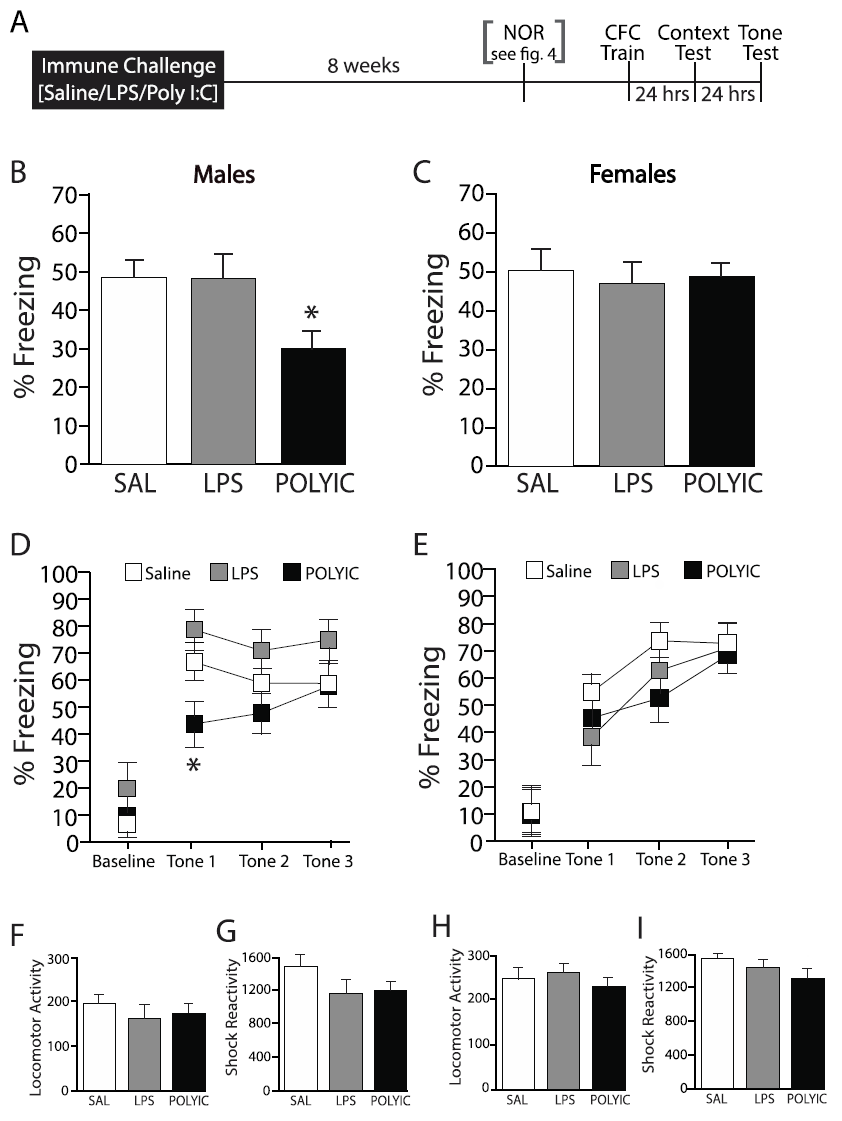
Context‐ and tone‐ fear conditioning was impaired in males long after subchronic Poly I:C. (A) Experimental timeline. Context fear conditioning was conducted in animals previously tested in novel object recognition (results in Fig. 4). (B) Freezing to the training context in males is decreased after subchronic Poly I:C, but not LPS treatment (n = 8-9 per group). (C) Context fear conditioning was not impaired in females after LPS or Poly I:C (n = 9 per group). (D) In males, Poly I:C, but not LPS, resulted in a mild impairment in tone fear conditioning. (E) Females showed no disruption of cued-fear conditioning after either immune challenge. (F,H) Locomotor activity and (G,I) shock reactivity during training was not altered after LPS or Poly I:C either sex. \**p* < 0.05 *cf* Saline

**Figure 7.**
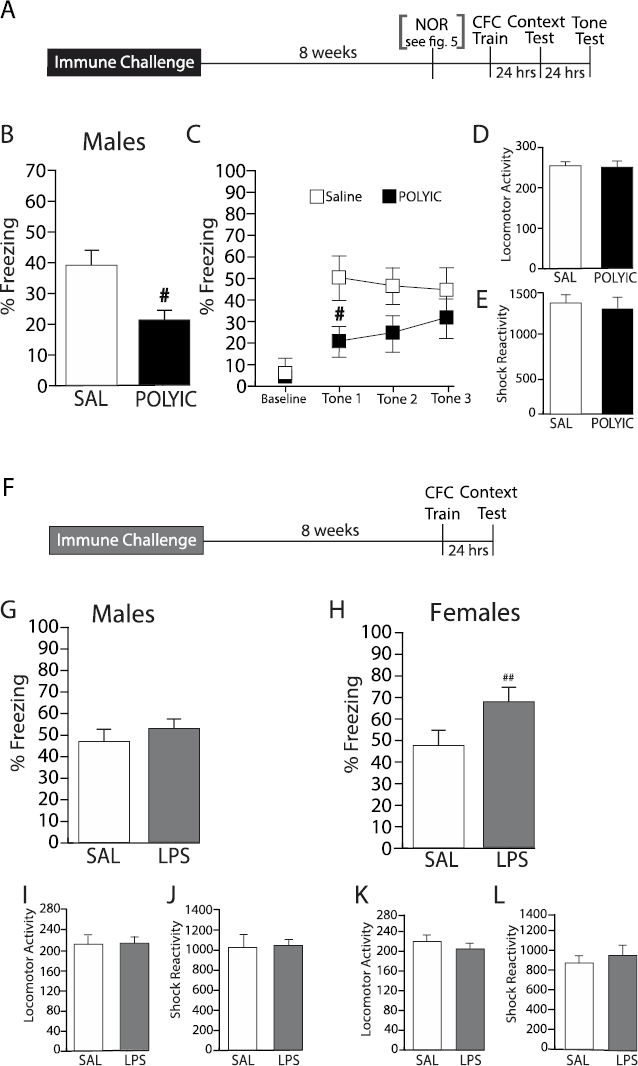
Replication: Disruption of fear conditioning eight weeks after subchronic Poly I:C, but not LPS, challenge. (A) Experimental timeline. Context fear conditioning was conducted in animals previously tested in 3-hour novel object recognition (results in Fig. 5). (B) Context fear conditioning and (C) tone fear conditioning was decreased in males eight weeks after subchronic Poly I:C challenge. (D,E) In context fear conditioning, locomotor activity and reactivity to the shock was not altered by subchronic Poly I:C. # *p*<0.01 *cf* saline. (F) Experimental timeline. Context fear conditioning was conducted in behaviorally naïve mice 8 weeks after LPS injections. (G,H) Foreground context fear conditioning was intact in both sexes. (I,K) Locomotor activity and (J,L) shock reactivity were not different between groups. # *p* < 0.05 *cf* saline, ##*p* < 0.01 *cf* saline.

#### 2.4.1 Novel Object Recognition.

The testing arena consisted of two rectangular opaque white chambers, (LWD: 40cm × 32cm × 32.5cm; 45 lux at center). Two novel object recognition protocols were used: (1) a hippocampal-dependent protocol where memory persists for at least 24 hours (Vogel-Ciernia & Wood, 2015); and (2) a more commonly assessed protocol that typicall results in short-term novel object recognition in which animals are tested 3 hours after training (Ballaz, Akil, & Watson, 2007).

To assess long-term novel object recognition memory, mice were first habituated to testing chambers (10 mins/day for 6 days). Mice received two 10-minute training trials spaced 3 hours apart in which they explored two identical objects. A single test session occurred 24 hours after the first training session, in which mice were replaced in the arena with one familiar object (from training) and one novel object (Vogel-Ciernia & Wood, 2015). Novel and familiar objects were counterbalanced across animals. The time spent exploring each object (animal’s nose within 2 cm of object) was measured automatically (Ethovision XT 9.0 tracking software; Ballaz, Akil, & Watson, 2007) and corroborated by an experimenter blind to experimental conditions. Novel object preference was calculated as the percent time spent at the novel object {100*[(Time exploring novel object)/(time spent exploring both objects)]}.

Short-term novel object recognition was assessed as a single, 10-minute training session followed 3 hours later by a test session as described above (Ballaz et al., 2007).

##### Data Analysis and statistics.

Habituation was analyzed using repeated measures (Day × ImmuneChallenge) ANOVA. Separate repeated measures (ObjExpl × ImmuneChallenge) ANOVA were conducted for each sex (e.g., Darcet et al., 2014). Planned comparisons were used (with LSD) to examine comparisons between exploration of novel and familiar objects for each group.

#### 2.4.2 Context Fear Conditioning.

The context fear conditioning apparatus consisted of rectangular chambers (LWD: 9.75″ × 12.75″ × 9.75″) containing grid floor rods connected to a shock generator, an enclosed sound-attenuating system, and a NIR camera (VID-CAM-MONO-2A) and Video Freeze software for automatic scoring of freezing behavior (MedAssociates, VT). During background context fear conditioning, mice were placed in the training context (rectangular box with white walls, lights on, an evenly sized grid floor, 70% ethanol odor) for 3-min, after which a 30-sec tone (10 kHz, 75 dB SPL) was presented co-terminating with a 2-sec 0.8mA footshock (Tronson et al., 2010). Mice were returned to their home cages immediately following training. We assessed automatically recorded locomotor activity during training to determine whether subchronic immune challenge alters movement or exploration prior to the shock (locomotor activity) (Cunningham & Sanderson, 2008) and locomotor response during the 2-sec shock (shock reactivity) to identify whether prior immune challenge resulted in differences in sensitivity to the aversive US (Tronson et al., 2010).

Twenty-four hours later, context fear memory was assessed. Mice were replaced in the training context for 3 minutes and freezing behavior was measured (Keiser et al., 2017). The following day, mice were tested for fear conditioning to the tone in a novel context (black angled walls, house lights off, staggered grid floors, 1% acetic acid odor). After 90 seconds in the novel context, three 30-sec tones separated by 60-sec intertrial intervals were presented. Freezing to the training context and to the tones were automatically scored using Video Freeze software (MedAssociates) (Anagnostaras et al., 2010; Keiser et al., 2017).

To assess the effect of subchronic immune challenge on late-occurring impairments in memory retrieval, animals were trained on background context fear conditioning one week after immune challenge and re-tested for context‐ and tone‐ fear memory seven weeks later. Foreground context fear conditioning was conducted as above, without presentation of the tone or tone fear tests (Keiser et al., 2017).

##### Data Analysis and Statistics.

Separate one-way ANOVA were used to assess the effect of prior immune challenge on context fear conditioning for each sex, and repeated measures ANOVA (Tone × ImmuneChallenge) were used to assess freezing to the cue in background fear conditioning. Post hoc tests (with LSD) were used to further assess specific group differences. Because males and females were always tested separately, group differences were compared within sex.

### 2.5 Immunohistochemistry.

Animals (n = 4 / group) were anesthesized (Avertin 480 mg/kg, i.p.) and transcardially perfused at 12 weeks post last immune challenge with 4% paraformaldehyde. 40 μM sections through the hippocampus were incubated with rabbit anti-mouse Iba-1 (1:10,000, WAKO), goat anti-mouse secondary (1:200, Vector Labs), and DAB chromagen (Sigma Aldrich, St Louis, MO). ImageJ (NIH, Bethesda, MD) was count microglia, and a fractal analysis plugin in ImageJ was used to determine microglial morphology [FracLac V. 2.5, (Karperien, Ahammer, & Jelinek, 2013)]. Microglial activation state is commonly assessed using number and morphological changes where cells shift from a resting ramified state with extended, branching processes, to an activated ameboid state with a larger cell body and retraction of processes (Kondo, Kohsaka, & Okabe, 2011; Singer et al., 2016; Weberpals et al., 2009). Fractal analysis is a sensitive and quantitative method to assess small changes in microglial shape using measures of pattern complexity and self-similarity (fractal dimension) as well as heterogeneity (lacunarity) (Karperien et al., 2013; Young & Morrison, 2018). An experimenter blind to the treatment conditions performed microglial counts and morphology analyses in the dentate gyrus molecular layer and CA1.

#### Data Analysis and Statistics.

Two-way ANOVA (Sex × ImmuneChallenge) were used to assess the effects of prior immune challenge on microglial activation for both microglia number and morphology. Post-hoc tests were used to further assess specific group differences.

### 2.6 Blood-Brain Barrier Permeability.

Blood-brain barrier permeability was assessed in male and female mice (n = 3) using the most sensitive method to detect disruption, sodium fluorescein (Birngruber et al., 2013; Kaya & Ahishali, 2011; Saunders et al., 2008). Sodium fluorescein (2%, i.p.) was injected 20 mins prior to blood collection and transcardiac perfusion with saline. Brain permeability index (BPI) was calculated comparing fluorescence (relative fluorescence units, RFU) in brain to fluorescence in serum [BPI = (RFU brain/brain weight)/(RFU Serum/serum volume)], and normalized to brain permeability index in control animals (Devraj, Guérit, Macas, & Reiss, 2018). We also used fluorescent microscopy to visualize qualitative differences in sodium fluorescein penetration in brain sections across multiple brain regions (20μM) (Nikon A1 laser scanning microscope (Devraj et al., 2018; Nikolian et al., 2018).

#### Data analysis and Statistics.

Independent sample t-tests were used to compare brain permeability index for Poly I:C‐ and saline-treated animals.

## 3. RESULTS

### 3.1 Systemic effects of Poly I:C and LPS.

To assess acute and long-lasting systemic effects of subchronic Poly I:C and LPS, we measured changes in weight over the eight week post-challenge period (Fig. 1). All mice gained weight across the 15 day treatment period (Injection: Males: *F*(4,88) = 43.01, *p* < 0.001; Females: *F*(4,96) = 42.72, *p* < 0.001). Across all 5 injections, male and female mice treated with LPS or Poly I:C showed decreased weight the day after injection (Males: ImmuneChallenge × Cycle: *F*(4,44) = 4.32, *p* < 0.01; LPS: *p* < 0.01; Poly I:C *p* < 0.05 vs injection day; Females: ImmuneChallenge × Cycle: *F*(4,48) = 4.63, *p* < 0.01; LPS *p* <0.05, Poly I:C *p* < 0.01 vs injection day Fig. 1B,C). For male mice treated with LPS, this weight loss persisted on the second day after injection (LPS: *p* < 0.05; Poly I:C *p* = 0.40 vs injection day). Saline-treated males showed no change in weight on average across the 2 days after injection (*p* = 0.22 and *p* = 0.52, respectively vs injection day) and saline treated females showed a small but significant weight gain (*p* < 0.01, *p* = 0.01, respectively vs injection day). Across the eight week time period, mice gained weight at equivalent rates regardless of prior treatment (Males: ImmuneChallenge × Day: *F*(2,22) = 1.93; *p* = 0.17; Females: *F*(2,22) = 1.91, *p* = 0.17).

### 3.2 Females, but not males, show object recognition memory deficits soon after subchronic immune challenge.

Novel object recognition memory was tested one week after Poly I:C using a paradigm resulting in a hippocampal-dependent memory 24 hours after training (Vogel-Ciernia & Wood, 2015; Fig. 2A). In males, we observed preference for exploration of the novel compared to the familiar object in both saline‐ and Poly I:C-treated groups (ObjExpl*: F*(1,14) = 7.82, *p* < 0.05) and no differences between treatments (ObjExpl × ImmuneChallenge: *F*(1,14) < 1; Fig. 2B). Therefore, males showed no deficit in object recognition one week after immune challenge. In contrast, subchronic immune challenge caused impairments of novel object recognition in females (ObjExpl × ImmuneChallenge: *F*(1,14) = 7.96, *p* < 0.05; Fig. 2C). Whereas saline-treated mice showed significantly greater exploration of the novel vs the familiar object (*p* < 0.01), Poly I:C-treated mice showed no such preference (*p* = 0.46). Thus females, but not males, are sensitive to impairments in memory in the first weeks after a subchronic immune challenge.

These memory deficits were not due to changes in locomotor activity or exploration. There were no differences in habituation to the testing chambers (Males: Day *F*(5,70) = 5.75; *p* < 0.01; Day × Drug: *F*(5,70) < 1; Females: Day *F*(5,70) = 5.75, *p* < 0.01; Day × Drug *F*(4,48) = 1.07, *p* = 0.37; Fig. 2D,F). All groups also showed similar locomotor activity (Males: *t*(14) <1; Females: *t*(14) < 1) and object exploration during training (all *t* < 1; Fig. 2E,G). No decreases in body weights were observed in Poly I:C-treated mice prior to habituation one week or two weeks after the last injection, prior to novel object recognition training (Males: ImmuneChallenge × Day: *F*(2,28) <1; Females: *F*(2,28) < 1).

### 3.3 Context fear conditioning is intact soon after immune challenge.

We next examined whether context‐ or tone-dependent fear conditioning was modulated one week after subchronic immune challenge, and if remote memory or retrieval processes were impacted seven weeks later (Fig. 3A). In males, we found no memory deficits after training one week after immune challenge, or during the remote memory test seven weeks later. Contrary to expectations, we observed that males showed an increase in freezing to context one week after Poly I:C (ImmuneChallenge *F*(1,13) = 5.17, *p* < 0.05; Time × Immune Challenge: *F*(1,13) = 2.19, *p* = 0.16; Time: *F*(1,13) < 1; 1week: *p* < 0.01; 8 weeks: *p* = 0.67; Fig. 3B). Subchronic immune challenge did not result in either memory deficits at one week, or in retrieval deficits eight weeks after the final injection.

In females, there were no effects of prior Poly I:C on context fear conditioning one week after subchronic immune challenge (ImmuneChallenge: *F*(1,14) = 2.1, *p* = 0.17; Time × ImmuneChallenge: *F*(1,14) < 1; Fig. 3C). However females in both groups showed a dramatic decrease in freezing to the context seven weeks after training (Time: *F*(1,14) = 27.30, *p* < 0.001; Fig. 3C), suggesting a decrease in memory retrieval, independent of prior immune challenge, at a remote time point in females.

We also observed no effect of Poly I:C on cued fear conditioning or retrieval one week or eight weeks after the final injection in either sex (Males: ImmuneChallenge: *F*(1,13) < 1; ImmuneChallenge × Time *F*(1,13) < 1; ImmuneChallenge × Time × Tone *F*(2,26) < 1; Fig. 3D; Females: ImmuneChallenge: *F*(1,14) = 3.32, *p* = 0.09; ImmuneChallenge × Time × Tone *F*(2,28) < 1; Fig. 3E) demonstrating that Poly I:C does not cause short-lasting changes in auditory fear conditioning or long lasting changes in amygdala-dependent fear memory retrieval.

Importantly, there were no differences in locomotor activity (Males: *t*(13) = 1.13, *p* = 0.28; Females: *t*(13) < 1; Fig. 3F,H) or response to the footshock during training (Males: *t*(13) = 1.57, *p* = 0.14; Females: *t*(13) < 1; Fig. 3G,I).

### 3.4 Deficits in 24-hour novel object recognition memory months after subchronic immune challenge in both males and females.

Novel object recognition memory was tested eight weeks after LPS or Poly I:C using a memory paradigm testing hippocampal-dependent memory 24 hours after training (Vogel-Ciernia & Wood, 2015; Fig. 4A). During test, we observed a significant detrimental effect of prior immune challenge on exploration of the novel object compared to the familiar object in males (ObjExpl × ImmuneChallenge: *F*(2,22) = 3.90, *p* < 0.05), with saline (*p* < 0.01), but not LPS (*p* = 0.76) nor Poly I:C (*p* = 0.93) treated males showing intact object recognition memory (Fig. 4B). In females, we observed a trend towards decreased novel object recognition after immune challenge (ObjExpl × ImmuneChallenge: *F*(2,14) = 2.97, *p* = 0.07). Importantly, only saline-treated females showed significantly more exploration of the novel object than the familiar object (saline: *p* < 0.05; LPS: *p* = 0.93; Poly I:C *p* = 0.46; Fig. 4C). Prior immune challenge disrupted novel object recognition in both males and in females.

There were no differences in locomotor activity nor habituation to the testing chamber for either sex (Males: Day: *F*(5,110) = 3.95, *p* < 0.01; Day × Drug: *F*(10,110) = 1.29, *p* = 0.27; Females: Day *F*(5, 120) = 21.17, *p* < 0.001; Day × Drug: *F*(10,120) < 1; Fig. 4D,F). During training, there were no differences in locomotor activity (Males: Drug: *F*(2,22) < 1; Females: *F*(2,24) < 1), and all animals showed similar exploration of objects, regardless of prior treatment (Males: *F*(2,24) < 1; Females *F*(2,26) < 1; Fig. 4E,G). Prior subchronic LPS or Poly I:C treatment did not affect locomotor activity, habituation to a new arena, or object exploration in males or females.

### 3.5 Disruption of 3-hour novel object recognition months after subchronic Poly I:C or LPS.

To further examine memory deficits eight weeks after subchronic Poly I:C in males, we used a short-term novel object recognition paradigm (Ennaceur, Neave, & Aggleton, 1997) that is less dependent on hippocampus. There were no differences between groups in object exploration (*t*(14) = 1.19, *p* = 0.25; Fig. 5C) nor locomotor activity (*t*(14) <1; Fig. 5D) during training. At test, novel object preference was significantly impaired in the Poly I:C-treated compared with the saline-treated mice (ObjExpl × ImmuneChallenge: *F*(1,14) = 21.55, *p* < 0.01; Fig. 5B), with significantly greater novel object exploration after saline (*p* < 0.05) and significantly lower novel object exploration after Poly I:C (*p* < 0.01).

Similarly, short-term novel object recognition was impaired in both sexes eight weeks after subchronic LPS challenge compared with saline-treated controls (Males: ObjExpl × ImmuneChallenge *F*(1,14) = 8.03, *p* < 0.05; Females: (ObjExpl × ImmuneChallenge *F*(1,14) = 32.88, *p* < 0.01; Fig. 5E,H). Only saline-treated animals showed preference for the novel object (Males: *p* < 0.05; Females *p* < 0.01), and LPS-treated animals showed no preference (Males: *p* = 0.23). Prior LPS had no effect on object exploration (Males: *t*(14)= 1.04, *p* = 0.31; Females: *t*(14) = 1.47, *p* = 0.17; Fig. 5F,I) or locomotor activity during training (all *t*(14) <1; Fig. 5G,J). Together, the data from both short‐ (Fig. 5) and long-term (Fig. 4) memory paradigms, demonstrate that novel object recognition memory is robustly impaired several months after either subhronic Poly I:C or LPS challenge.

### 3.6 Impaired context and tone-dependent fear conditioning months after Poly I:C challenge in males but not females.

We next tested context‐ and tone‐ fear conditioning. Given prior testing in these animals, fear conditioning took place approximately 10 weeks after the subchronic immune challenge (Fig. 6A). In males, prior Poly I:C but not LPS challenge caused deficits in context fear conditioning (*F*(2,24) = 3.63, *p* < 0.05; Poly I:C *p* < 0.05, LPS: *p* = 0.97 cf saline; Fig. 6B). In contrast, females showed no deficits in context fear conditioning after either LPS or Poly I:C (*F*(2,24) < 1; Fig. 6C).

In males, fear conditioning to the tone was also impaired, with lower freezing to the first test tone in Poly I:C-treated males (ImmuneChallenge: *F*(2,22) = 4.87, *p* < 0.05; Tone × ImmuneChallenge: *F*(4,44) = 2.77, *p* < 0.05; Tone: *F*(2,44) = 1.50 *p* = 0.24; *p* < 0.05 cf saline*; p* < 0.01 cf LPS; Fig. 6D). There were no differences between groups in freezing to the novel context (*F*(2,22) = 2.15*, p* = 0.14). In contrast, females showed no alterations of tone fear conditioning after either immune challenge (ImmuneChallenge: *F*(2,44) = 1.31, *p* = 0.29; Tone × ImmuneChallenge *F*(4,48) = 1.38; *p* = 0.25; Tone: *F*(2,48) = 19.48, *p* < 0.001 Fig. 6E), with no differences in freezing to the novel context (*F*(2,24) < 1; Fig. 6E).

Neither locomotor activity during training (Males: *F*(2,24) <1, Females: *F*(2,24) < 1; Fig. 6F,H) nor response to shock (Males: *F*(2,22) = 2.08, *p* = 0.15, Females: *F*(2,24) = 1.22, *p* = 0.32; Fig. 6G,I) differed across groups in either sex, thus deficits in context fear conditioning were not due to a failure to detect the shock US. Together, these findings demonstrate that males, but not females, are sensitive to disruption of fear conditioning long after subchronic Poly I:C challenge.

It is particularly striking that the same female animals showed deficits in hippocampal-dependent novel object recognition (Fig. 4C) but not context fear conditioning (Fig. 6C), which also depends on hippocampus. Similarly, after LPS, the same males showed impairments of novel object recognition (Fig. 4B) but not context fear conditioning (Fig. 6B).

### 3.7 Replication: Poly I:C, but not LPS disrupts fear conditioning months after subchronic challenge.

Due to the surprisingly long-lasting nature of context fear conditioning after subchronic Poly I:C challenge in males, we next determined whether this finding was robust and replicable. Once again, we observed that Poly I:C induced disruption of both context‐ and tone‐ fear conditioning after subchronic immune challenge in males. During test, mice displayed significantly lower freezing in the training context compared with saline controls (*t*(14) = 2.75, *p* < 0.05; Fig. 7B). Tone fear conditioning was once again impaired after Poly I:C (ImmuneChallenge *F*(1,14) = 6.67, *p* < 0.05; Tone: *F*(2, 28), *F* <1; ImmuneChallenge × Tone F(2,28) = 1.27, *p* = 0.23), with no differences in freezing to the novel context (*t*(14) = 1.15, *p* = 0.27; Fig. 7C). There were no observed changes in locomotor activity (*t*(14) < 1; Fig. 7D) or reactivity to the shock (*t*(14) < 1; Fig. 7E) in Poly I:C-treated mice.

In a separate experiment, we replicated the lack of effect of subchronic LPS on persistent memory deficits in both sexes. Because the previous experiment was conducted several weeks after the 8-week timepoint at which we observed novel object recognition deficits (Fig. 6A), we tested the possibility that LPS-induced memory deficits recover more quickly than those induced by Poly I:C by testing animals at eight weeks after subchronic LPS challenge (Fig. 7F). We replicated the finding that subchronic LPS caused no disruption in context fear conditioning in either sex eight weeks after challenge (ImmuneChallenge *F*(1,28) = 3.56, *p* = 0.07; Sex *F*(1,28) < 1; ImmuneChallenge × Sex *F*(1,28) < 1; Fig. 7G,H). There were no observed changes in locomotor activity or reactivity to the shock (all F< 1).

### 3.8 Subchronic Poly I:C challenge does not induce sustained hippocampal microglia activation or increased blood brain barrier permeability.

We observed no differences in microglia number (CA1: all *F*(1,12) < 1; Fig. 8A-B; DG: ImmuneChallenge: *F*(1,12) = 1.94, *p* = 0.19, Sex × ImmuneChallenge: *F*(1,12) < 1; Fig. 8D-E) or morphology (CA1: fractal dimension: ImmuneChallenge: *F*(1,12) = 2.63, *p* = 0.13, Sex × ImmuneChallenge: *F*(1,12) < 1, lacunarity: ImmuneChallenge: *F*(1,12) = 1.78, *p* = 0.21, Sex × ImmuneChallenge: *F*(1,12) < 1, Fig. 8C-D; DG: fractal dimension: all F(1,12) < 1, lacunarity ImmuneChallenge: *F*(1,12) = 2.16, *p* = 0.17, Sex × ImmuneChallenge: *F*(1,12) < 1, Fig. 8G-H) in the dorsal hippocampus months after Poly I:C challenge. Similarly, blood-brain barrier showed no increase in permeability long after Poly I:C treatment. Poly I:C did not alter the brain permeability index (ImmuneChallenge: *t*(1,12) = 1.06, *p* = 0.31; Fig. 8I) and there was no increases in sodium fluorescein stainingof saline and Poly I:C treated males or females across any brain region assessed (data not shown).

**Figure 8.**
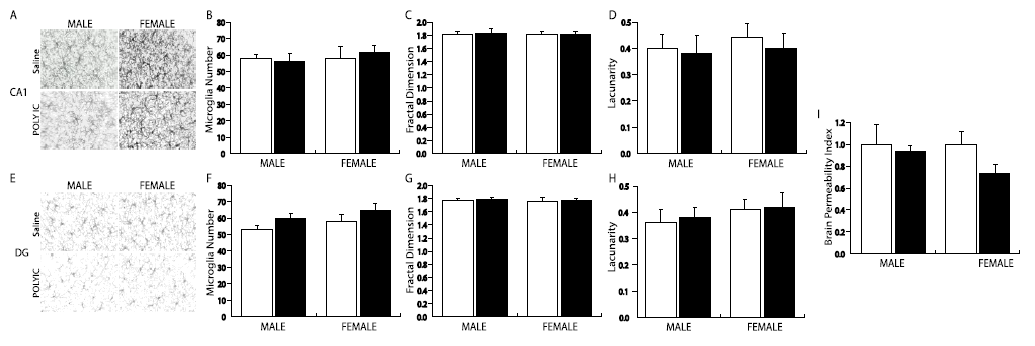
Subchronic Poly I:C did not induce sustained microglial activation or blood-brain barrier permeability. (A,B) Representative images microglia (40x) in males and females in CA1 and dentate (molecular layer) of dorsal hippocampus. (C,D) Microglial numbers are unaltered in the CA1 or dentate gyrus (molecular layer) months after Poly I:C challenge. (E-H) Prior Poly I:C did not alter microglial morphology, as measured by decreased self-similarity (fractal dimension) and increased heterogeneity (lacunarity). (I) Prior Poly I:C did induce sustained blood-brain permeability.

## 4. DISCUSSION

These findings are the first to demonstrate sex-specific patterns of memory deficits after a subchronic, systemic immune challenge. Object recognition memory was impaired in both sexes eight weeks after immune challenge, but females also exhibited early deficits on novel object recognition. Furthermore, males but not females showed delayed deficits in context and tone-dependent fear conditioning after Poly I:C. Retrieval of an established memory was unaffected. Thus, prior immune challenge specifically caused dysregulation of memory formation. Importantly, memory deficits were independent of sustained sickness or neuroimmune activation as indicated by normal weight gain, microglial activation, and blood-brain barrier function. Together, these findings demonstrate progressive changes in neural function and sex-specific patterns of memory deficits emerging over the weeks and months following a mild, subchronic immune challenge. The delayed emergence of memory deficits in the absence of ongoing overt microglial activation suggests that the transient immune response exerts lasting indirect effects on neural function (Tchessalova, Posillico, & Tronson, 2018).

In males, both LPS and Poly I:C disrupted object recognition, however only Poly I:C resulted in deficits in context and auditory fear conditioning. This suggests that specific patterns of immune activation lead to different patterns of neural dysfunction. Indeed, distinct types of immune responses are observed in various animal models of inflammation (Heremans, Dillen, Dijkmans, Grau, & Billiau, 1989). Here, LPS and Poly I:C bind to different toll-like receptors (TLR4 and TLR3, respectively), recruit different signaling pathways, and thereby exert differential effects on the brain (Doyle et al., 2003). Acutely, both LPS and Poly I:C cause memory impairments (Cloutier, Kavaliers, & Ossenkopp, 2012), but LPS causes greater decreases in wheel running and locomotor activity (Hopwood, Maswanganyi, & Harden, 2009), and Poly I:C causes more deleterious effects in developmental models (Arsenault, St-Amour, Cisbani, Rousseau, & Cicchetti, 2014). We observed that prior Poly I:C, but not LPS, disrupted fear conditioning, suggesting that viral mimetics induce a greater long-lasting impact on fear-related circuitry, including amygdala. We chose doses of LPS and Poly I:C based on similar effects on behavior and weight loss over the two-week injection period and similar doses to those previously compared (e.g., Arsenault et al., 2014). It is possible, however, that differences in the intensity or duration of immune activation caused the broader impairments in memory after Poly I:C compared with LPS. Indeed, tolerance to repeated injections is commonly observed with LPS but not to Poly I:C (Soszynski, Kozak, & Szewczenko, 1991), suggesting that in our experiments, animals treated with Poly I:C may have experienced more chronic neuroimmune activity than those treated with LPS. Nevertheless, given the differences in patterns of cells (Starkhammar et al., 2012), cytokines (Kimura et al., 2004), and downstream effectors (Suh, Zhao, Derico, Choi, & Lee, 2013) activated by LPS and Poly I:C, directly comparing levels of immune activation remains difficult. Identifying how each kind of immune challenge (e.g., polymicrobial sepsis, gram-negative and gram-positive bacteria, viral mimics, injury/surgery/heart attack, etc.) targets different neural structures and functions will be critical for understanding dysregulation of cognitive and affective processes that persist long after a transient immune event.

In contrast to the delayed memory deficits in males, females showed greater vulnerability to memory impairments in the first week after the immune challenge. This pattern of early and persistent deficits in females in contrast to delayed, broader memory impairments in males is consistent with recent findings from our lab showing more rapid activation and resolution of neuroinflammatory signaling after systemic immune challenge in females compared with males (Speirs & Tronson, 2018). These findings are consistent with sex differences in peripheral immune responses (Furman et al., 2014; Klein & Flanagan, 2016) and differential regulation of neuroimmune activation (Acaz-Fonseca et al, Arevalo, 2015; Bodhankar et al, 2015; Santos-Galindo et al, 2011). Sex differences in susceptibility to memory dysfunction may therefore be due to differences in pattern, time course, or strength of immune activation.

That males, but not females show deficits in fear conditioning after a subchronic immune challenge also suggest sex differences in susceptibility to immune modulation across brain regions. This is consistent with previous studies of sex differences in neuroimmune activity. Floor example, after chronic stress, males but not females showed ongoing microglial activation in basolateral amygdala (Bollinger, Collins, Patel, & Wellman, 2017). Alternatively, sex differences in strategy and mechanisms of fear memory formation may mediate differential susceptibility of males to disruption by prior Poly I:C (Keiser et al., 2017; Shansky, 2018). Fear conditioning triggers sex-specific patterns of activation across brain regions (Keiser et al., 2017; Lebron-Milad et al., 2012) and signaling mechanisms (Keiser & Tronson, 2015; Mizuno & Giese, 2010). Thus male-specific disruption of fear conditioning may be due to sex differences in neuroimmune signaling, and/or to disruption of circuits or mechanisms required for fear conditioning in males but not females.

This model of subchronic immune activation is an important new tool for identifying the neural mechanisms underlying progressive cognitive deficits in the weeks and months after recovery from illness. Long-lasting memory deficits have previously been described after serious illness in humans, and in models of sepsis including high-dose LPS stimulation or cecal ligation and puncture (Huerta et al., 2016; Ming et al., 2015). Here we demonstrated that repeated administration of a lower dose of LPS or Poly I:C is sufficient to cause delayed and persistent memory deficits without overt, persistent neuroimmune activation. The lack of microglial activity observed here is notable since previous work using high doses of LPS or cecal ligation and puncture that have observed memory deficits have also observed persistent microglial activation within the hippocampus (Kondo et al., 2011; Singer et al., 2016; Weberpals et al., 2009). Our findings suggest that microglia activity, and the active neuroimmune state they indicate, are not necessary for enduring or emerging memory deficits after immune challenge. This emphasizes the importance of identifying the impact of transient neuroimmune signaling on neuronal signaling and neural function.

Although we did not observe microglial activation, it is likely that more subtle neuroimmune changes persist long after a subchronic immune challenge (Tchessalova et al., 2018). For example, persistent microglial priming persists to adulthood after early life immune challenge (Hoeijmakers, Heinen, van Dam, Lucassen, & Korosi, 2016; Williamson & Bilbo, 2013). In astrocytes, states changes are observed during aging (Clarke et al., 2018), and astrocytic markers are increased for least 30 days after immune challenge or surgery, even in the absence of increased cytokine production (Fu et al., 2014; Jin, Feng, Feng, Lu, & Qi, 2014). Such altered reactivity of neuroimmune cells significantly impact responses to subsequent immune challenge or stressor in adulthood and during aging (Norden, Muccigrosso, & Godbout, 2015; Schwarz & Bilbo, 2011) but neither are commonly regarded as ongoing immune activation *per se*. Nevertheless, together with recent data demonstrating the critical role of astrocytes in memory processes and synaptic plasticity (Adamsky et al., 2018; Gao, Suzuki, Magistretti, Lengacher, & Pollonini, 2016; Suzuki et al., 2011), findings suggest the possibility that enduring changes in astrocyte function may mediate the emergence of memory deficits observed in this study.

Additionally, previous studies have identified several neuronal processes that are persistently dysregulated after a transient immune challenge. For example, a single high dose of LPS results in persistent alterations of cholinergic function (Ming et al., 2015), impairments in neurogenesis (Ormerod et al., 2013), epigenetic modifications (Schaafsma et al., 2015), and late-occurring striatal neurodegeneration (Liu et al., 2008; Qin et al., 2007). Other studies have demonstrated decreases in spine density in hippocampus (Chavan et al, 2012; Volpe et al, 2015) and amygdala (Huerta et al., 2016) one month after such as cecal ligation and puncture. These findings, together with the data described here, suggest that neuroimmune activation as a consequence of systemic immune challenge has long-term impact on neuronal function, even after overt microglial activation has resolved. Future work is needed to delineate specific mechanisms that mediate the delayed and enduring memory deficits observed here.

Rodent models that use exogenous stimuli including LPS and Poly I:C have limitations in their ability to mimic human illnesses. For example, although high doses of LPS is commonly used in animal models of sepsis, there are concerns about differences in immune response to endotoxin in humans versus rodent models limiting the efficacy of this model (Fink, 2014; Rittirsch, Hoesel, & Ward, 2007). In neuroscience research, Poly I:C has been predominantly used to model maternal viral infection and maternal immune activation on the effects of neural development (Reisinger et al., 2015). Neither LPS nor Poly I:C accurately model the course of bacterial or viral infection, or the full time-course of an illness. Nevertheless, these are important tools for triggering transient and robust immune – and neuroimmune – activation, and thereby useful models for identifying both short‐ and long-term changes in cognitive and affective processes, neural function, and their underlying molecular mechanisms in the absence of ongoing disease processes.

Importantly, this model is sensitive to sex differences in vulnerability to and time course of emergent memory deficits, and differential effects of bacterial‐ and viral-like immune triggers. These findings thus provide a basis from which to identify immune and non-immune mechanisms that drive memory deficits long after recovery from illness, surgery, or injury. As such, this model will be an important tool for distinguishing the direct immune mechanisms and indirect impact on neural function that contribute to early and progressive memory impairments after neuroimmune activation in both sexes.

These findings demonstrate that subchronic systemic inflammation causes sex-specific patterns of memory decline over the following months. Both males and females are susceptible to impairments in memory after immune challenge, but females are more vulnerable to early-onset memory deficits, whereas males are more susceptible to later development of impairments across multiple memory systems. Surprisingly, these memory impairments are not associated with persistent neuroimmune mechanisms. Additional studies will be required to identify the long-lasting neural changes that mediate such delayed and persistent memory impairments. This subchronic immune challenge model is thus a valuable tool for identifying how systemic inflammation initiates memory decline and memory-related disorders. Understanding the persistent neural changes as a consequence of transient neuroimmune activation will be critical for development of strategies to prevent cognitive decline after major illness or chronic inflammation in women and in men.

**FUNDING AND DISCLOSURES:** This work was funded by NIH/NIMH R00 MH093459 to NCT. The authors declare no conflicts of interest.

## ACKNOWLEDGEMENTS

We would like to thank Dr. Anuska Andjekovic-Zochowska and Dr. Svetlana Stamatovic for their expertise and assistance with blood brain barrier permeability assays; and Dr. Ashley Keiser, Caitlin Posillico, and Brynne Raines for their helpful comments on this manuscript.

## CONTRIBUTIONS

D.T. and N.C.T. designed the experiments. D.T. ran experiments and collected the data. D.T. and N.C.T. analyzed the data, wrote the manuscript, and revised the paper.

